# TIP60/KAT5-mediated regulation of apoptotic sensitivity during DNA damage-induced senescence

**DOI:** 10.64898/2026.11.12.766234

**Authors:** Shiksha Dutta, Nicolas Malaquin, Erwan Goy, Audrey Carrier-Leclerc, Guillaume B Cardin, Francis Rodier

## Abstract

Cellular senescence and apoptosis are two distinct outcomes of DNA damage. It is unclear what determines the fate decision and the characteristics of senescent cells. Chromatin modifiers are likely candidates, as they can influence the environment for repair of DNA double-strand breaks as well as the transcription of senescent genes. Here we report that the lysine acetyltransferase TIP60/KAT5 is a dosage-sensitive determinant of senescent cell identity in normal human fibroblasts. Although TIP60 is not required for the senescence-associated proliferation arrest of cells after a single dose of 10 Gy of ionising radiation, the resolution of 53BP1 and γH2AX foci as well as the amplitude of the senescence-associated secretory phenotype (SASP) as measured by transcript and protein levels of secreted factors is dependent on the levels of TIP60. Furthermore, the sensitivity of senescent cells to apoptosis is also modulated by TIP60 levels. Cells with low levels of TIP60 show reduced p53 output and high levels of the anti-apoptotic protein MCL-1 and are completely resistant to the senolytic ABT-263. Cells with high levels of TIP60 are modestly more sensitive to ABT-263. These results place TIP60 upstream of the divergence of senescent phenotypes and establish that proliferation arrest and the SASP are uncoupled by the level of a single chromatin modifier. We further identify TIP60 as a candidate determinant of senolytic / senomorphic response and suggest that MCL-1 co-inhibition could be used as a rational addition in contexts where TIP60 levels are low.

## Introduction

Cell fate decisions after genotoxic insults unfold through a multistep molecular signaling process initiated by DNA double-strand breaks (DSBs), leading to the deployment of the DNA damage response (DDR) ^1,2^. The outcome of this signalling cascade depends on the extent and persistence of damage but also on pre-existing cell states, for example, the levels of anti-apoptotic proteins ^3^. When DNA damage is repairable, transient DDR signalling allows re-entry into the cell cycle, whereas extensive or irreparable damage commits cells to cell-cycle arrest, senescence or apoptosis ^4–6^. How this life-or-death decision is taken remains unclear but involves interactions between DDR activation and pre-existing cell state or identity. At the molecular level, DSBs trigger the immediate local recruitment of the MRE11–RAD50–NBS1 (MRN) complex, which senses the break and recruits the ataxia-telangiectasia mutated (ATM) kinase ^7^. ATM activation is marked by autophosphorylation at serine 1981 (pS1981-ATM)^8^, after which ATM phosphorylates histone H2AX on serine 139 (γH2AX) across the chromatin surrounding DSBs ^9^. This mark recruits repair factors such as 53BP1 and BRCA1, which direct repair pathway choice^10^, while ATM signals through CHK2 and the p53-p21 axis to activate cell cycle checkpoints and/or pro-apoptotic factors to determine cell fate ^11,12^.

Cellular senescence is often triggered by persistent DNA damage and defined by a stable senescence-associated (SA) proliferation arrest (SAPA) that prevents the expansion of cells at risk of neoplastic transformation^13,14^. Senescent cells also display increased SA-β-galactosidase (SA-β-Gal)^15^ activity and develop an SA secretory phenotype (SASP), comprising cytokines, chemokines, growth factors and proteases that remodel the tissue microenvironment^16,17^. Although the SASP supports wound repair and embryonic development, a chronic SASP drives inflammation, tumour progression and age-related disease^18^. The SASP highly depends on the transcription factor nuclear factor kappa-light-chain-enhancer of activated B cells (NF-κB)^19^, which acts together with associated transcription factors C/EBPβ^20^ and GATA4^21^ to drive inflammatory gene expression. Its amplitude is further set by ATP-dependent chromatin remodelling complexes, including SWI/SNF^22^ and NuA4/TIP60^23^. Additional chromatin regulators such as HMGB2^24^, macroH2A1^25^ and lamin B1^26^ shape the chromatin landscape at SASP gene loci. The SASP is also amplified by cyclic GMP–AMP synthase (cGAS) and stimulator of interferon genes (STING), which sense cytoplasmic DNA from chromatin fragments released into the cytoplasm of senescent cells ^27^. Interestingly, although ATM is essential for the initiation and maintenance of the SASP, its early canonical activation within the DDR is not sufficient to trigger the SASP^28^, which is rather associated with recruitment of the MRN-ATM activation complex on the chromatin and independent from ATM kinase activity ^5,29^. Overall, multiple chromatin-associated activities appear linked to the activation of the NF-κB SASP, suggesting that chromatin remodellers could be key players in regulating SASP dynamics during cell fate decisions^18^.

In contrast to senescence, apoptosis is a programmed form of cell death that eliminates damaged cells without inducing inflammation ^30,31^. The tumour suppressor p53 lies at the centre of the decision between these fates, transactivating either cell-cycle arrest genes such as *CDKN1A* (p21) or pro-apoptotic genes such as *BBC3* (PUMA), *PMAIP1* (NOXA) and *BAX* ^32,33^. The choice between these transcriptional programmes depends on the post-translational modification status of p53, the availability of its cofactors, the abundance of anti-apoptotic proteins such as myeloid cell leukaemia-1 (MCL-1), and the cellular context ^34,35^. Accordingly, phosphorylation of p53 at serine 46 and acetylation at lysine 120 (K120), together with cofactors such as ASPP1 and ASPP2, direct p53 towards pro-apoptotic targets ^36,37,38,39^. Induction of these pro-apoptotic targets triggers mitochondrial outer membrane permeabilisation (MOMP), cytochrome *c* release and the caspase cascade leading to cell death ^40^.

By definition, senescence (survival) and apoptosis (death) are mutually exclusive ^41^. However, once established, senescent cells further resist apoptosis, in part by shifting the balance of the B-cell lymphoma 2 (BCL-2) family towards its anti-apoptotic members ^42^. This dependence constitutes a therapeutic vulnerability, as BCL-2 family inhibitors such as ABT-263 (navitoclax) selectively eliminate senescent cells ^43^. Accordingly, such senolytic agents have been proposed for combination with genotoxic therapy in a “one-two punch” approach, whereby senescent cells generated by the primary therapy are subsequently cleared ^44^. However, whether any apoptotic signal translates into cell death is not determined solely by p53 activity but by the pre-existing balance of pro- and anti-apoptotic BCL-2 family proteins at the mitochondria, a state termed apoptotic priming ^3^. The degree of priming is intrinsic to each cell type, as demonstrated by lymphoid cells, which are highly primed and undergo rapid p53-dependent apoptosis following irradiation, and fibroblasts, which are poorly primed and preferentially engage cell-cycle arrest ^45^. Although the mechanisms that establish this apoptotic identity remain poorly understood, its stable and cell-type-specific nature resonates with some role for chromatin regulators in its determination ^37^, similar to their established contribution to the senescence programme ^22^.

A key regulator at the interface of chromatin and the DDR is the lysine acetyltransferase KAT5 (also known as Tat-interactive protein, 60 kDa, TIP60), the catalytic subunit of the NuA4/TIP60 complex ^46^. TIP60 is recruited to DSBs in an MRN-dependent manner, where binding to histone H3 trimethylated at lysine 9 (H3K9me3) stimulates its acetyltransferase activity towards ATM ^47,48^. TIP60 further acetylates histones H4 and H2A to limit 53BP1 binding and favour homologous recombination (HR) ^49^. It also acetylates p53 at K120, a modification required for transcription of PUMA and BAX ^37,38^. Beyond the DDR, TIP60 is a haploinsufficient tumour suppressor and connects to two arms of SASP regulation, acting as a cofactor for the NF-κB subunit RelA/p65 and acetylating cGAS ^50–52^. This acetyltransferase activity is not confined to sites of damage. The NuA4/TIP60 complex functions as a transcriptional co-activator at active promoters^53,54^, and proliferation is dependent on this function. Conditional deletion of KAT5 in primary fibroblasts arrests cells without inducing p53, INK4A or ARF ^55^. Transcription factors that drive the G1/S transition recruit the complex to their target promoters, E2F1 to late G1 target promoters^56^ and MYC to chromatin occupied by MYC^57^. TIP60 therefore integrates chromatin state, ATM signalling and p53-dependent cell fate. Despite these connections, whether TIP60 actively modulates cell fate decisions and senescence after the infliction of persistent DNA lesions remains undefined.

Here, we investigate the impact of TIP60 on the establishment and maintenance of SA phenotypes and cell fate. Using irradiation (IR) to initiate damage-induced senescence synchronously in normal human cells and following the DDR over time, we identify TIP60 as a determinant of the rate at which damaged cells resolve persistent DNA damage foci and of the phenotypic characteristics of the senescent state they subsequently adopt. TIP60 is dispensable for SAPA but rate-limiting for the SASP, demonstrating that the two defining features of senescence can be uncoupled by the level of a single chromatin modifier. This uncoupling extends to cell fate, as TIP60 levels set the apoptotic priming of senescent cells and thereby their susceptibility to senolytic elimination. Together, our findings place TIP60 at the chromatin-level interface between the DDR and cell fate and establish the SASP and senolytic sensitivity as tunable features rather than fixed properties of the senescent state, for better therapeutic understanding.

## Materials and Methods

### Cell line and culture conditions

Primary human BJ foreskin fibroblasts obtained from J. Smith (University of Texas, San Antonio) served as the parental cell line for this study. Cells were maintained at ambient oxygen in Dulbecco’s Modified Eagle’s Medium (DMEM) supplemented with 10% fetal bovine serum (FBS), 2.5 µg/ml Fungizone (amphotericin B), 100 U/ml penicillin and 100 µg/ml streptomycin. Unless stated otherwise, early-passage fibroblasts (<35 cumulative population doublings, PD) were used. PD was calculated at each passage as PD = previous PD + log₂ (cells harvested / cells seeded). For conditioned media (CM) experiments, we washed dishes three times with low-serum medium (0.1% FBS) and cultured cells in the same medium for up to 24 h. In all other experiments, the medium was replaced every 2 days. Lentiviral particles were produced in 293FT packaging cells (Invitrogen) as described previously ^29^. All cell lines used in this study were periodically tested for mycoplasma contamination using a Mycoplasma PCR Detection Kit (Abcam) and were confirmed to be negative.

### Viruses and infections

Viruses were produced as described previously ^29,58^. Virus titres were adjusted to infect 95–99% of cells in a final volume of 1 ml in a T25 flask with 4 µg/ml polybrene. To label nuclei, parental BJ fibroblasts were first transduced with lentivirus encoding TurboGFP (green fluorescent protein) fused to a C-terminal SV40 nuclear localisation signal (tGFP-NLS) (pLenti CMV Blast DEST, 706-1, Addgene #17451) ^58^. GFP-positive cells were isolated by fluorescence-activated cell sorting (FACS) rather than by blasticidin selection and are referred to as BJ-tGFP-NLS. All genetically modified lines used in this study were subsequently derived from this BJ-tGFP-NLS line by lentiviral transduction. For TIP60 overexpression, BJ-tGFP-NLS cells were transduced with lentivirus encoding untagged and HA-tagged TIP60 (pLenti CMV Hygro DEST, w117-1, Addgene #17454)^58^, with EGFP (pLenti CMV/TO Puro DEST, 670-1, Addgene #17293)^58^ as the corresponding control. For TIP60 depletion, cells were transduced with lentiviral pLKO.1-puro vectors carrying short hairpin RNAs (shRNAs) against TIP60 and GFP as a non-targeting control (originally Open Biosystems, now Horizon Discovery). All shRNA clone identifiers and target sequences used in the study are provided in Supplementary Table S1. Infections were followed 48 h later by selection with puromycin (1 µg/ml) or hygromycin B (100 µg/ml), as appropriate for the respective vector, for six days. Stable cells were either used directly or stored at -80 °C.

### Irradiation

Cells were exposed to a single 10 Gy dose of X-irradiation (IR) at a dose rate of approximately 3.3 Gy/min using a Gammacell 3000 Elan ¹³⁷Cs irradiator (Best Theratronics).

### Real-time imaging-based cell proliferation assay

Cells were seeded in 96-well plates at 1,500 cells per well. After 24 h, cells were either irradiated (10 Gy) or left untreated (control), and proliferation was monitored for 10 and 7 days, respectively. Plates were imaged in an IncuCyte S3 Live-Cell Imaging System (Sartorius) maintained at 37 °C and 5% CO₂. Two fields per well were captured every 6 h using a 10× objective in phase contrast and green fluorescence. Cell number was quantified with IncuCyte S3 software as the number of green-fluorescent objects per well. Objects were segmented using a fixed mask defined by a minimum green fluorescence intensity and minimum and maximum object areas, chosen to select the compact nuclear tGFP-NLS signal. Mask parameters were held constant across all lines and time points within a plate, and were defined separately for irradiated and non-irradiated plates to account for the differences in cell morphology and fluorescence intensity after IR. Relative cell number was calculated for each well by normalising the GFP object count at each time point to the count at day 0, defined as the time of IR or mock treatment, 24 h after seeding.

### Real-time cell death imaging and senolytic assay

Cell death was assessed using propidium iodide (PI), a cell-impermeant DNA dye that labels dead cells ^59^. PI (Sigma-Aldrich) was added to the culture medium at 0.5 µg/ml, and cells were imaged as described for the real-time proliferation assay. The percentage of dead cells was quantified with IncuCyte S3 software as the number of PI-positive red nuclei relative to the total number of tGFP-NLS–labelled green nuclei at each time point. To assess senolytic sensitivity, cells were treated with 1 µM navitoclax (ABT-263; APExBIO) and an equivalent volume of DMSO (0.1% final concentration) in medium containing 0.5 µg/ml PI. Treatment began on day 9 post IR for irradiated cells and six days after the end of antibiotic selection for untreated cells. Plates were imaged every 4 h over the subsequent 24 h, and the percentage of dead cells was quantified at each time point as described above.

### Reverse transcription quantitative PCR (RT-qPCR)

Cells were seeded in 100-mm dishes and irradiated (10 Gy) 24 h later or left untreated. RNA was harvested 3 days (untreated controls) or 9 days (irradiated) post IR. Total RNA was extracted using the RNeasy Mini Kit (Qiagen) and stored at −80 °C. One microgram of total RNA was reverse transcribed using the SuperScript III First-Strand Synthesis System (Invitrogen, 200 U) according to the manufacturer’s instructions. The resulting cDNA was diluted 1:10, and 1 µl was used per reaction with sequence-specific primers (400 nM) and SYBR Select Master Mix (Applied Biosystems, Thermo Fisher Scientific). Primers were designed using NCBI Primer-BLAST (http://blast.ncbi.nlm.nih.gov/) and are listed in Supplementary Table S2. Reactions were run on a StepOnePlus Real-Time PCR System (Applied Biosystems) (UDG activation at 50 °C for 2 min; AmpliTaq activation and denaturation at 95 °C for 2 min; then 40 cycles of 95 °C for 15 s, 60 °C for 1 min and 72 °C for 30 s). Data were analysed using StepOne software (Applied Biosystems). Gene expression was normalised to TATA-binding protein (TBP) gene expression, and relative expression was calculated using the 2^-ΔΔCt method.

### EdU incorporation assay

Cells were cultured in 8-well Falcon™ chambered cell culture slides (Corning). After 24 h, cells were either irradiated (10 Gy) or left untreated. EdU (5-ethynyl-2′-deoxyuridine; 10 µM, Invitrogen) was added to the medium for 48 h pulses spanning days 1–3, 5–7 and 7–9 post IR. Untreated cells were pulsed over days 5–7, 12–14 and 19–21 following 6 days of antibiotic selection. Cells were washed three times with TBS and fixed in 10% neutral-buffered formalin (Sigma) for 10 min at room temperature. EdU incorporation was detected using the Click-iT protocol with Alexa Fluor 647 dye (Invitrogen). Slides were mounted with ProLong Gold Antifade Mountant containing DAPI (Thermo Fisher Scientific). Images were acquired on a Zeiss AxioObserver Z1 fluorescence microscope (Carl Zeiss) using a 20× objective with AxioVision software (Carl Zeiss). The percentage of EdU-positive nuclei was quantified in >150 nuclei in three different fields per condition using ImageJ ^60^.

### Immunofluorescence

Cells were seeded in 8-well Falcon™ chambered cell culture slides (Corning). At days 3, 5 and 7 post IR, cells were fixed in 10% neutral-buffered formalin (Sigma-Aldrich) for 10 min at room temperature, washed in PBS and permeabilised in PBS–0.25% Triton X-100 (Sigma-Aldrich) for 10 min. Slides were washed in PBS, and non-specific binding was blocked for 1 h in PBS containing 1% bovine serum albumin (IgG-free, protease-free; Jackson ImmunoResearch) and 4% normal donkey serum (Sigma-Aldrich) (blocking buffer). Primary antibodies against γH2AX (mouse monoclonal, JBW301, Merck Millipore) and 53BP1 (rabbit polyclonal, NB100-305, Novus Biologicals) were diluted 1:1000 in blocking buffer and incubated overnight at 4 °C. Slides were washed three times in PBS–0.05% Tween 20 (Thermo Fisher Scientific) and incubated with secondary antibodies diluted 1:5000 in blocking buffer for 1 h at room temperature. After three further washes in PBS–0.05% Tween 20, slides were mounted and imaged as described for the EdU incorporation assay. Nuclei were identified by DAPI staining, and γH2AX and 53BP1 foci were counted in >150 nuclei in three different fields per condition using ImageJ ^60^.

### Senescence-associated β-galactosidase staining

Senescence-associated β-galactosidase (SA-β-gal) activity was assessed essentially as described ^15^. Irradiated cells were stained at day 9 post IR; untreated cells were stained at days 7, 14 and 21 following the 6-day antibiotic selection. Cells were washed once with PBS, fixed in 10% formalin (Sigma-Aldrich) for 5 min at room temperature and washed again with PBS. Cells were then incubated for 16 h at 37 °C in staining solution containing 1 mg/ml X-gal (5-bromo-4-chloro-3-indolyl-β-D-galactopyranoside, from a 20 mg/ml stock in dimethylformamide), 5 mM potassium ferricyanide, 5 mM potassium ferrocyanide, 150 mM NaCl, 2 mM MgCl₂ and 40 mM citric acid/sodium phosphate buffer at pH 6.0. Cells were then washed twice with PBS, and nuclei were counterstained with 300 nM DAPI in PBS for 5 min at room temperature, followed by two washes in PBS. Five randomly selected fields per well were imaged on an EVOS FL Digital Inverted Microscope (Thermo Fisher Scientific) using a 10× objective with brightfield and DAPI channels. The percentage of SA-β-gal-positive cells was calculated as the number of blue-stained cells divided by the total number of DAPI-stained nuclei in the same field.

### Analysis of SASP factor secretion in conditioned medium

Cells were seeded in 96-well plates (Corning) and irradiated (10 Gy) 24 h later or left untreated. Conditioned medium (CM) was prepared essentially as described previously.^61^ Briefly, cells were washed three times with low-serum medium (0.1% FBS) and incubated in the same medium for 16 h, at day 9 post IR for irradiated cells and at days 7, 14 and 21 following the 6-day antibiotic selection for untreated cells. Cell numbers were determined by counting tGFP-NLS-positive nuclei on the IncuCyte S3 Live-Cell Imaging System (Sartorius) at the time of collection. IL-6 was quantified using the Human IL-6 Quantikine ELISA Kit (R&D Systems) according to the manufacturer’s instructions. Values were normalised to cell number and expressed as fold change in secreted protein relative to the corresponding control. CM was also analysed for 40 secreted factors by multiplex electrochemiluminescence immunoassay using the V-PLEX Human Biomarker 40-Plex Kit (Meso Scale Discovery) according to the manufacturer’s instructions. Concentrations of secreted protein per millilitre were normalised to cell number and expressed as log₂ fold change relative to the corresponding control.

### Statistics

Unless stated otherwise, all data are presented as the mean ± standard deviation of three independent experiments. Statistical analyses were performed using GraphPad Prism 11.0 Software (GraphPad Inc.). Experiments comparing cell lines at a single endpoint were analysed by one-way ANOVA, and experiments with two variables (cell line and time point) by two-way ANOVA. In all cases, ANOVA was followed by Dunnett’s multiple comparisons test against the relevant control line (EGFP or shGFP.3). Differences were considered statistically significant at p < 0.05, and significance is indicated in the figures as *p < 0.05, **p < 0.01, ***p < 0.001 and ****p < 0.0001.

## Results

### TIP60 dosage modulates proliferation without altering cell fate

TIP60 behaves as a haploinsufficient tumour suppressor^50^, suggesting that its cellular functions depend on its abundance. We therefore manipulated TIP60 expression in normal human BJ fibroblasts, a well-established model of cell senescence.^58^ For depletion, we used lentiviruses to stably deliver two independent shRNAs of different efficiency, shTIP60.4 and shTIP60.5, which reduced *KAT5* mRNA to approximately 40% and 20%, respectively, compared to the level in non-targeting shGFP.3 control cells (Fig. 1A). For overexpression, we similarly stably delivered untagged TIP60 and HA-tagged TIP60 (TIP60HA) using lentiviruses raising *KAT5* mRNA above the levels of EGFP control cells (Fig. 1B).

**Figure 1.**
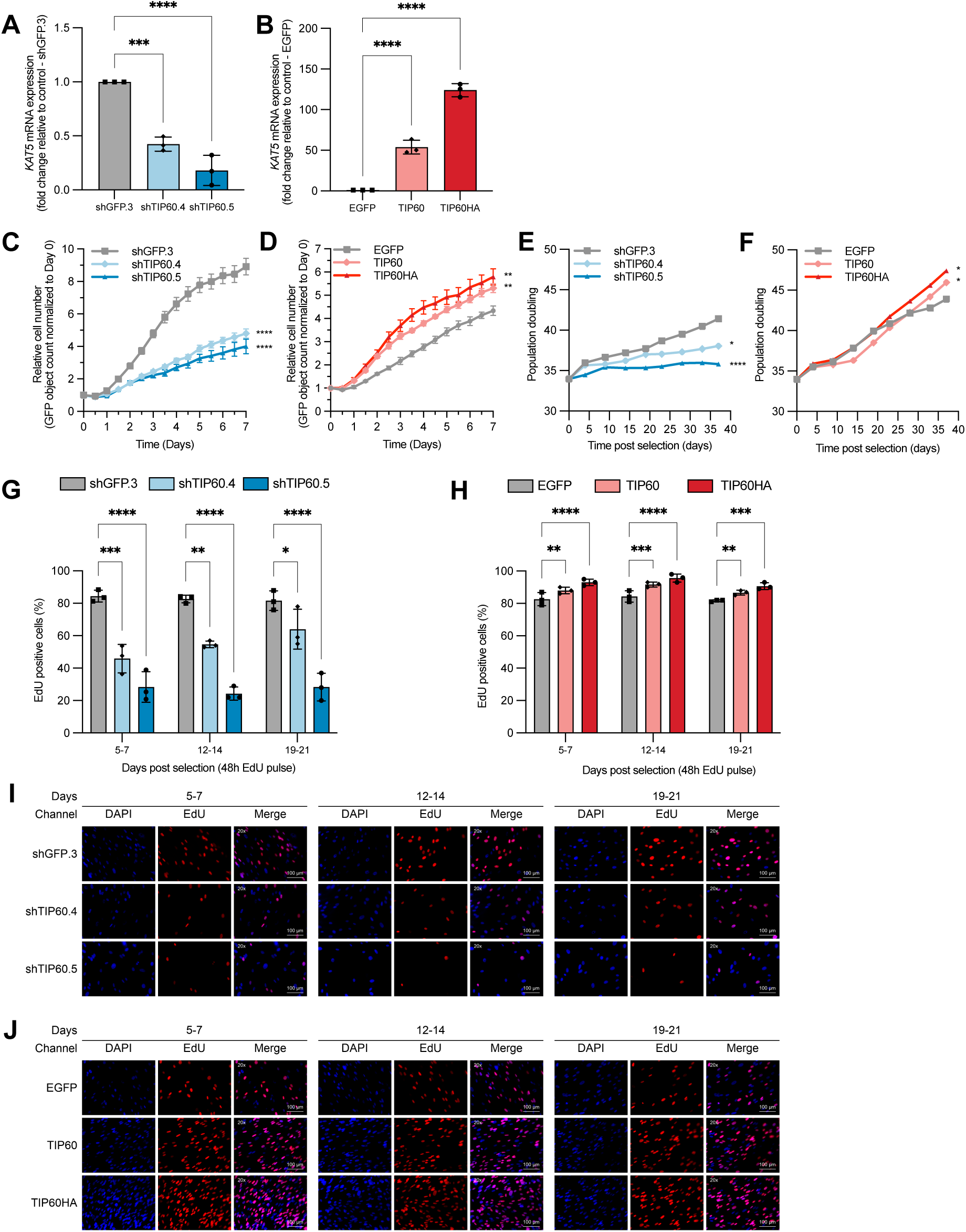
TIP60 dosage modulates proliferation without altering cell fate. (A, B) *KAT5* mRNA levels measured by RT-qPCR in untreated BJ-tGFP-NLS fibroblasts expressing (A) a non-targeting shRNA (shGFP.3) or one of two shRNAs against TIP60 (shTIP60.4, shTIP60.5), or (B) expressing EGFP, untagged TIP60 or HA-tagged TIP60 (TIP60HA). RNA was harvested 3 days after seeding. Expression was normalised to *TBP* and is shown as fold change relative to the respective controls (shGFP.3 or EGFP) using the 2^–ΔΔCt method. Mean ± SD of three independent experiments, one-way ANOVA followed by Dunnett’s multiple comparisons test against the control line. (C, D) Real-time proliferation of (C) knockdown and (D) overexpressing lines. Cells were seeded at 1,500 per well in 96-well plates and imaged every 6 h for 7 days on an IncuCyte S3 system. Cell number was quantified as the number of tGFP-NLS-positive nuclei per well and normalised to the count at day 0, defined as 24 h after seeding. (E, F) Cumulative population doublings (PD) of (E) knockdown and (F) overexpressing lines during serial passaging after the end of antibiotic selection. PD was calculated at each passage as previous PD + log₂(cells harvested/cells seeded). (G, H) Percentage of EdU-positive nuclei in (G) knockdown and (H) overexpressing lines after 48-h EdU pulses applied on days 5–7, 12–14 and 19–21 post 6-day selection. EdU was detected by Click-iT chemistry with Alexa Fluor 647 and nuclei were counterstained with DAPI. More than 150 nuclei in three fields were scored per condition. Mean ± SD from three independent experiments. (I, J) Representative images of the EdU incorporation assay for (I) knockdown and (J) overexpressing lines at each pulse window, showing DAPI (blue), EdU (red) and the merged channels. Images were acquired with a 20× objective. Scale bar, 100 µm. C–H were analysed by two-way ANOVA followed by Dunnett’s multiple comparisons test against the respective control line; in C–F, one representative of four independent experiments is shown: mean ± SD of 6 replicate wells. Asterisks indicate the comparison at the final time point. *p < 0.05, **p < 0.05, ***p < 0.005, ****p < 0.0005.

Proliferation arrest is among the primary hallmarks of senescence,^13,14^ and apoptotic sensitivity can be judged relative to baseline survival.^41^ We first defined the effect of TIP60 level variations on these parameters before introducing genotoxic stress. Real-time imaging of tGFP-NLS nuclei over 7 days showed that TIP60 depletion slowed the expansion of undamaged cultures relative to controls, and that the effect was stronger in shTIP60.5 than in shTIP60.4, in keeping with the degree of knockdown (Fig. 1C, S1A). Slower expansion was not caused by cell loss, as imaging with propidium iodide (PI), which labels dead cells,^59^ detected few PI-positive cells over the same period (Fig. S1A, S1B). Overexpression had the opposite effect. Cultures proliferated faster than controls, with the larger gain in the line expressing more TIP60 (Fig. 1D, S1D), again without any significant change in cell death (Fig. S1D, S1E). Therefore, TIP60 level alone is sufficient to shift the growth rate of undamaged fibroblasts in accordance with expression levels.

To assess whether this difference was maintained in the long term, cells were passaged serially for five weeks after selection. Control cells divided steadily throughout. shTIP60.4 cells accumulated population doublings (PD) at a slower rate, whereas shTIP60.5 cells expanded only during the first two weeks and then reached a plateau, after which the population showed negligible proliferation (Fig. 1E; reproduced in two independent experiments started at different cumulative PD (Fig. S1C)). The pattern was reversed in response to TIP60 overexpression; cells accumulated PD more rapidly than controls, with higher-expressing TIP60HA cells consistently the furthest ahead (Fig. 1F; reproduced in two independent experiments (Fig. S1F)). The effect of TIP60 level on proliferation is therefore stable over many population doublings.

We next measured active DNA synthesis at three windows after selection, covering the period over which the PD curves diverged. Every time, more than 80% of control nuclei showed active S-phase entry, whereas the fraction was lower in TIP60-depleted lines (Fig. 1G, 1I). Because a sustained decline in DNA synthesis could reflect premature senescence, SA-β-gal activity^15^ was scored at the same time and remained at background in all knockdown lines (Fig. S1G, S1I). Conversely, TIP60 overexpression produced a small but reproducible DNA synthesis increase, with TIP60HA cells showing the larger effect (Fig. 1H, 1J), while SA-β-gal activity was unchanged (Fig. S1H, S1J). The reduced population growth of TIP60-depleted culture therefore reflects a smaller proportion of cells entering DNA replication rather than cell death or senescence, while the modest growth advantage of overexpressing culture reflects a larger proportion of replicating cells. Together, these data show that TIP60 sets the basal proliferative state of normal fibroblasts in a dose-dependent manner while leaving viability and senescence unchanged.

### TIP60 promotes DNA repair and limits persistent DDR signalling after IR

TIP60-modulated cells were next exposed to a single dose of 10 Gy IR, which induces DSBs and senescence synchronously in normal human fibroblasts through persistent DDR signalling ^5,29^. *KAT5* mRNA was measured on day 9 post IR when cells are normally senescent and transcript levels remained reduced in TIP60-depleted cells (Fig. 2A) and elevated in overexpressing cells relative to IR-treated controls (Fig. 2B). We next followed proliferation after IR by real-time imaging over 10 days (Fig. 2C, 2D, S2A, S2B). Every line reached a plateau, including the TIP60-manipulated cells, showing that the IR-induced proliferation arrest occurs irrespective of TIP60 level. However, the lines differed in the extent of proliferation before reaching the plateau and arrest. Knockdown lines expanded less than controls (Fig. 2C, S2A), whereas overexpressing lines expanded more, with TIP60HA cells the furthest ahead (Fig. 2D, S2B).

**Figure 2.**
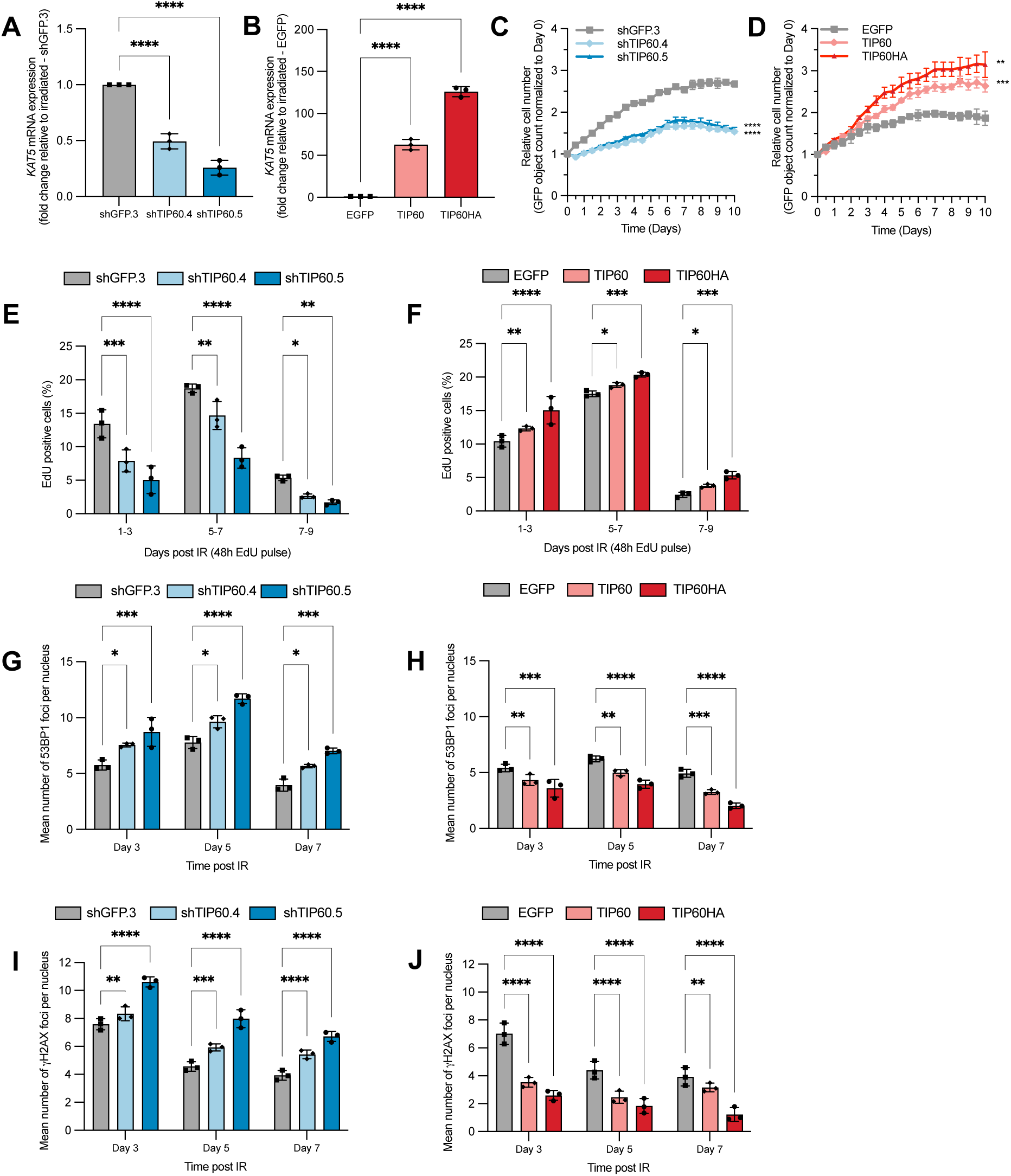
TIP60 promotes DNA repair and impairs persistent DDR signalling after IR. Cells were exposed to a single 10 Gy dose of IR. (A) Knockdown and (B) overexpressing lines, *KAT5* mRNA measured by RT-qPCR on day 9 post IR, normalised to *TBP* and expressed relative to IR-treated controls using the 2^–ΔΔCt method. (C) Knockdown and (D) overexpressing lines, showing proliferation after IR followed by real-time imaging of GFP-positive nuclei every 6 h for 10 days, with cell number normalised to the first time point. Cells were seeded at 1,500 per well, and data are from one representative of four independent experiments, with 6 replicate wells per line. Asterisks refer to the comparison at the final time point. (E) Knockdown and (F) overexpressing lines, proportion of cells synthesising DNA, analysed by EdU incorporation over 48 h windows spanning the acute stress, transition and senescent phases (days 1–3, 5–7 and 7–9 post IR). (G) Knockdown and (H) overexpressing lines, number of 53BP1 foci per nucleus, and (I) knockdown and (J) overexpressing lines, number of γH2AX foci per nucleus, analysed by immunofluorescence during the acute stress, transition and senescent phases. At least >150 nuclei were scored per condition and experiment. Data are the mean ± SD from three independent experiments. Statistical significance was assessed by one-way ANOVA (A, B) or two-way ANOVA (C–J) with Dunnett’s multiple comparisons test against controls. *P < 0.05, **P < 0.05, ***P < 0.005, ****P < 0.0005.

To confirm that the plateau reflected a loss of DNA synthesis, we examined the proportion of cells synthesising DNA over 48 h windows during the acute stress, transition and senescent phases (days 1–3, 5–7 and 7–9 post IR) (Fig. 2E, 2F, S2C, S2D). In contrast to more than 80% of non-irradiated cells that synthesised DNA over an equivalent window, the proportion in IR-treated controls was strongly reduced during the acute stress phase. During the transition phase, DNA synthesis recovered briefly, consistent with a subset of surviving cells attempting a final round of proliferation before the arrest, which accounts for the residual expansion of the growth curves before they reach their plateau (Fig. 2C, 2D). DNA synthesis then declined to near-complete loss in the senescent phase. Within each phase, however, the proportion of DNA synthesis was lower in shTIP60.4 and further reduced in shTIP60.5 cells (Fig. 2E, S2C), and higher in TIP60 and TIP60HA cells (Fig. 2F, S2D), reflecting observations in non-irradiated cells (Fig. 1G-1J). TIP60 level thus modulates the proportion of cells that complete residual cycles after damage but does not prevent them from entering a stable proliferation arrest.

Senescence after IR is sustained by persistent DNA damage foci that retain 53BP1 and γH2AX long after the initial lesions are formed.^5,29^ TIP60 is recruited to DSBs, where its acetyltransferase activity promotes repair by HR, in part by acetylating H4 and H2AX to open the chromatin around the break and by limiting 53BP1 retention at damaged chromatin.^46,48,49,62,63^ To test whether TIP60 abundance influences DNA damage foci, 53BP1 and γH2AX foci were assessed per nucleus. TIP60 depletion increased 53BP1 and γH2AX foci in every phase, with shTIP60.5 cells carrying the most and retaining them longest (Fig. 2G, 2I, S2E). Overexpression reduced both markers in every phase, with TIP60HA cells carrying the fewest and almost clearing γH2AX foci by the senescent phase (Fig. 2H, 2J, S2F). These results suggest that TIP60 abundance influences repair capacity, with elevated TIP60 driving resolution of DSBs and reduced TIP60 leaving a larger burden of unrepaired lesions that sustain senescence signalling.

### TIP60 modulates the amplitude of the senescence-associated secretome

Persistent DDR signalling and chromatin remodeling are involved in SASP regulation as the secretory program develops over several days after damage^5,16,28^. Having found that TIP60 level influences DNA damage foci persistence and senescence proliferation arrest, we investigated whether TIP60 also impact the secretory program of senescent cells. IL-6 secretion measured by ELISA at day 10 post IR in senescent cell populations was reduced in TIP60-depleted cells, with shTIP60.5 cells secreting the least (Fig. 3A), and elevated in overexpressing cells, with TIP60HA cells secreting the most (Fig. 3B). This dose relationship was also observed in non-irradiated cells, where baseline IL-6 secretion was lower in TIP60-depleted lines and higher in overexpressor at every point sampled over three weeks of culture after selection (Fig. S3A, S3B). To determine whether the TIP60 dependency extended beyond the SASP factor IL-6, we measured the mRNA expression of a panel of SASP genes at day 9 post IR. Nearly every transcript was reduced in shTIP60.5 relative to control cells (Fig. 3C) and increased in TIP60HA relative to control cells (Fig. 3D). The affected genes included IL-6, IL-8, IL-1α, IL-1β, GROα and GM-CSF, together with matrix metalloproteinases, IGFBPs, TGFB2 and VEGF, indicating that TIP60 can act at the level of SASP genes expression. To confirm that these transcriptional changes translated into secreted protein, CM was profiled by multiplex immunoassay across cytokine, chemokine and angiogenesis panels. Secreted factors were predominantly reduced in TIP60-depleted cells (Fig. 3E) and predominantly elevated in overexpressing cells (Fig. 3F). The same shifts in transcript and protein were present in baseline non-irradiated cells (Fig. S3C–S3F), albeit at much reduced levels, indicating that TIP60 levels scale the entire secretory programme rather than a subset of damage-responsive factors.

**Figure 3.**
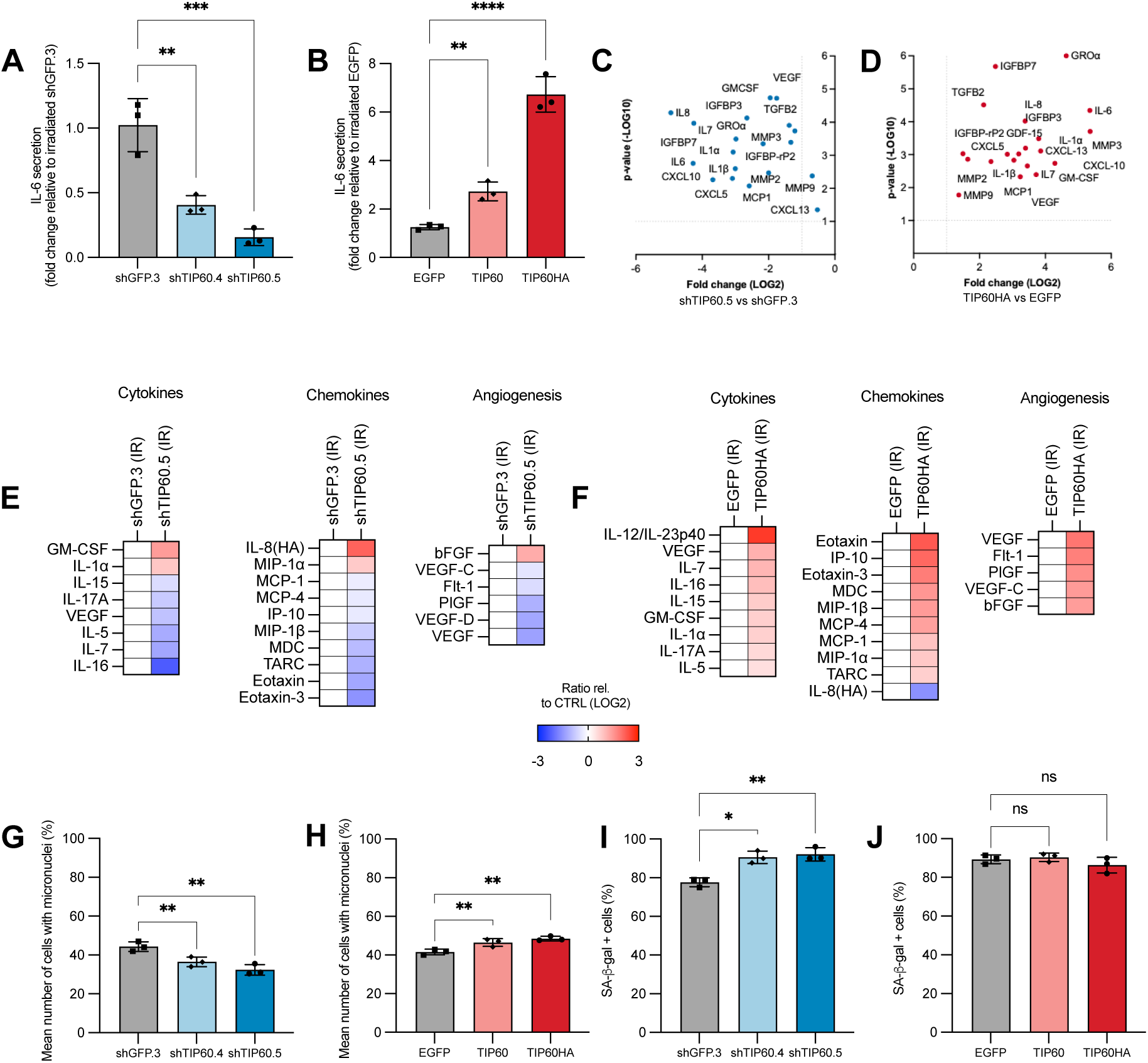
TIP60 dosage modulates the amplitude of the senescence-associated secretome. Cells were exposed to a single 10 Gy dose of IR. (A) Knockdown and (B) overexpressing lines, IL-6 secretion measured by ELISA in CM collected at day 10 post IR after 16 h of incubation, expressed as fold change relative to IR-treated shGFP.3 or EGFP controls. (C) Knockdown and (D) overexpressing lines: expression of SASP genes measured by RT-qPCR at day 9 post IR, shown as log2 fold change in (C) shTIP60.5 relative to shGFP.3 and in (D) TIP60HA relative to EGFP, with -log10 p-value. Normalisation and statistics as in Fig. 1A, 1B. (E) Knockdown and (F) overexpressing lines, secreted factors measured in CM collected on day 9 post IR by multiplex immunoassay (MSD V-PLEX cytokine, chemokine and angiogenesis panels), shown as log2 ratio, normalised to cell number, in (E) shTIP60.5 relative to shGFP.3 and in (F) TIP60HA relative to EGFP. IR denotes IR-treated cells. (G) Knockdown and (H) overexpressing lines, proportion of cells with micronuclei at day 7 post IR, scored in the DAPI channel of the images shown in Fig. S2C, S2D. At least >150 cells were scored per condition and experiment. (I) Knockdown and (J) overexpressing lines, proportion of SA-β-gal-positive cells at day 10 post IR. Data are mean ± SD of three independent experiments. Statistical significance was assessed by one-way ANOVA with Dunnett’s multiple comparisons test against controls. *P < 0.05, **P < 0.05, ***P < 0.005, ****P < 0.0005, ns not significant.

Senescent cells accumulate micronuclei, which arise when cells with unrepaired DNA pass through mitosis ^64,65^ and promote the SASP through cGAS-STING sensing of cytosolic chromatin and downstream NF-κB activation^27,66,67^. While TIP60 is related to both arms of this pathway via cGAS acetylation^52^ and as an NF-κB coactivator^68^, we also tested whether TIP60 influences genomic instability via accumulating micronuclei. We scored the proportion of cells with micronuclei at day 7 post IR, the end of the transition phase (Fig. S2C, S2D). TIP60 depletion reduced the proportion of cells carrying micronuclei (Fig. 3G), whereas overexpression increased it (Fig. 3H), in the same direction as the secretome and opposite to the DNA damage foci.

Finally, we asked whether the altered secretome reflected a change in the proportion of cells that became senescent. SA-β-gal staining^15^ at day 10 post IR was high in all lines (Fig. S3G, S3H). TIP60 depletion modestly increased the proportion of SA-β-gal-positive cells (Fig. 3I, S3G), consistent with the larger burden of persistent foci in these lines, whereas overexpression had no effect (Fig. 3J, S3H). The reduced secretome of TIP60-depleted cells therefore cannot be explained by fewer cells entering senescence, and the elevated secretome of overexpressing cells is not accompanied by more. Together, TIP60 sets the amplitude of the secretory phenotype of IR-induced senescent fibroblasts, with depletion dampening and overexpression amplifying the SASP.

### TIP60 modulates the apoptotic sensitivity of IR-induced senescent cells

Senescent cells resist apoptosis through elevated expression of anti-apoptotic BCL-2 family proteins, which allows them to persist despite sustained damage signalling^42^. This dependence makes them vulnerable to BCL-2 family inhibitors such as ABT-263, which selectively eliminate senescent cells^43,69^. TIP60 acetylates p53 and is required for the induction of its pro-apoptotic targets^37,38^. We therefore asked whether TIP60 level moulds the balance between survival and death factors in senescent cells. mRNA expression of an apoptosis-related gene panel was measured on day 9 post IR in the lines with the largest TIP60 perturbations. In shTIP60.5 cells, *p53* its arrest target *p21* and its pro-apoptotic targets *PUMA, NOXA, BAD* and *FAS* were reduced relative to controls, whereas *MCL-1*, an anti-apoptotic BCL-2 family member that is not targeted by ABT-263, was increased (Fig. 4A). In TIP60HA cells, *p53* and its targets were elevated together with *NOTCH1* and *BAX*, whereas the anti-apoptotic genes *BCL-2* and *MCL-1* were only marginally changed (Fig. 4B). TIP60 level thus shifts the transcriptional balance of senescent cells, with depletion favouring survival through reduced *p53* output and elevated *MCL-1*, and overexpression favouring apoptotic priming.

**Figure 4.**
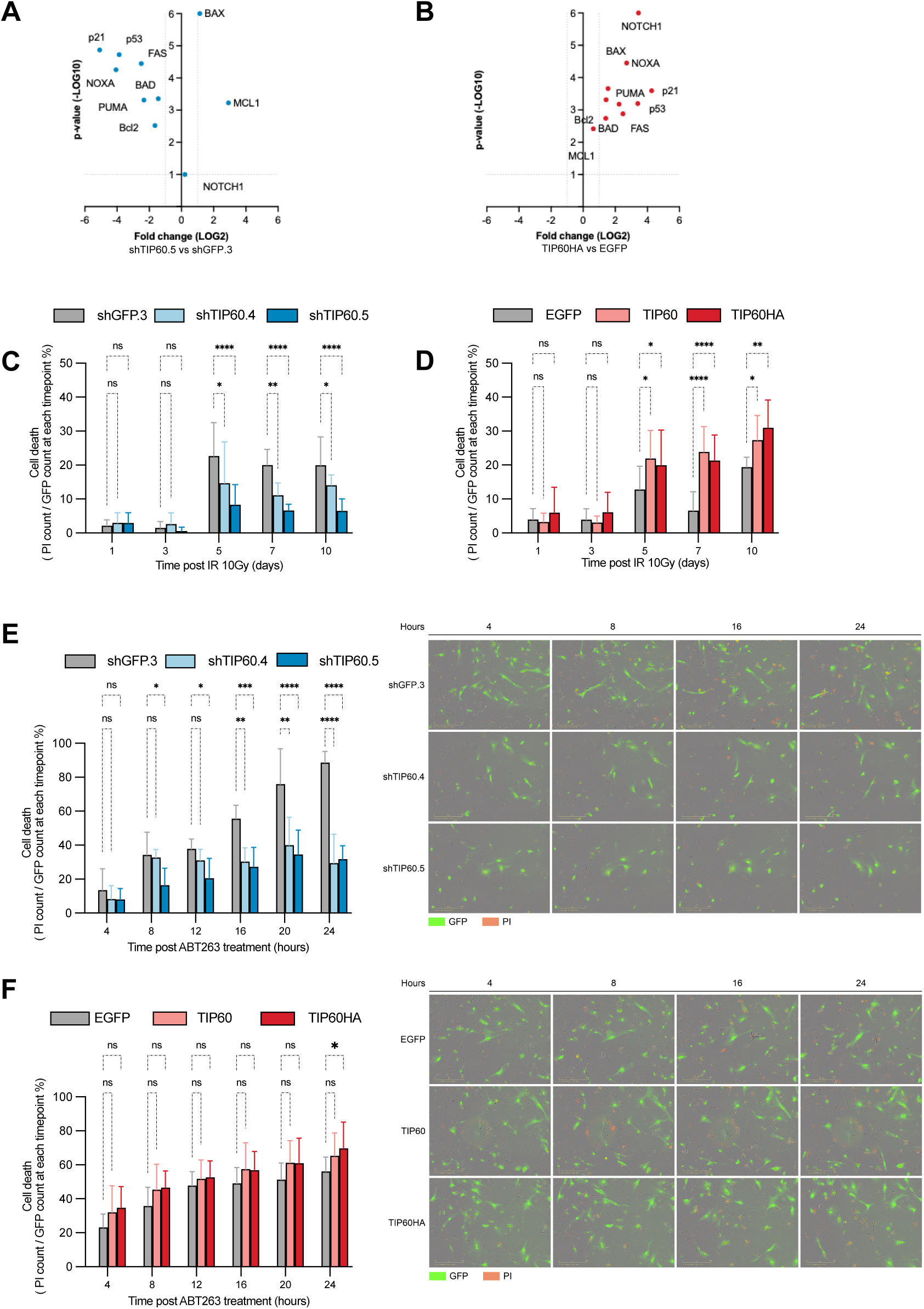
TIP60 modulates the apoptotic sensitivity of IR-induced senescent cells. Cells were exposed to a single 10 Gy dose of IR. (A) Knockdown and (B) overexpressing lines, expression of apoptosis-related genes measured by RT-qPCR on day 9 post IR, shown as log2 fold change in (A) shTIP60.5 relative to shGFP.3 and in (B) TIP60HA relative to EGFP with -log10 p-value. Normalisation and statistics as in Fig. 1A, 1B. (C) Knockdown and (D) overexpressing lines, cell death after IR measured by real-time imaging as the number of PI-positive cells relative to the number of GFP-positive nuclei at days 1, 3, 5, 7 and 10 post IR. (E) Knockdown and (F) overexpressing lines, cell death in IR-induced senescent cells treated with 1 µM ABT-263 at day 9 post IR, measured as in C and D every 4 h for 24 h. Representative images at 4, 8, 16 and 24 h post-treatment show GFP-positive nuclei (green) and PI-positive dead cells (red). Data are one representative of three independent experiments, mean ± SD of 3 technical replicates. Statistical significance was assessed by two-way ANOVA with Dunnett’s multiple comparisons test against shGFP.3 or EGFP controls at each time point. *P < 0.05, **P < 0.01, ***P < 0.001, ****P < 0.0001, ns not significant.

To determine whether this translated into cell death, we followed PI uptake by real-time imaging for 10 days after IR (Fig. 4C, 4D). Death was low and indistinguishable between lines during the acute stress phase (days 1–3 post IR). From the transition phase (days 5–7 post IR), death increased in controls and remained elevated through the senescent phase (days 7–9 post IR). In TIP60-depleted cells, this rise was diminished, with shTIP60.5 cells showing the least cell death (Fig. 4C, S2A), whereas in overexpressing cells it was enhanced (Fig. 4D, S2B). None of these differences were present in non-irradiated cells, in which death remained low and equal across lines for a week after selection (Fig. S1A, S1B, S1D, S1E). TIP60 level therefore determines the proportion of cells that die as they transition into senescence, without affecting the survival of proliferating cells.

Finally, we tested whether TIP60 level modulates the response of senescent cells to senolytic treatment. IR-induced senescent cells on day 9 post IR were treated with ABT-263, and death was followed by real-time imaging over 24 h (Fig. 4E, 4F). Control senescent cells died progressively, with most of the population PI-positive by 24 h. TIP60-depleted senescent cells were largely protected, with cell death in both knockdown lines plateauing within the first hours and remaining far below controls thereafter (Fig. 4E). Overexpressing senescent cells showed modestly more cell death than controls at every time point, reaching significance for TIP60HA at 24 h (Fig. 4F). Non-irradiated cells treated with ABT-263 at day 6 post seeding were insensitive to the drug regardless of TIP60 level (Fig. S4A, S4B), confirming that it acts selectively on the senescent state. Together, TIP60 modulates the apoptotic sensitivity of IR-induced senescent fibroblasts, with depletion conferring resistance to both spontaneous and senolytic-induced death through reduced p53-dependent priming and MCL-1 upregulation, and overexpression enhancing it.

## DISCUSSION

Our study identifies TIP60 as a dosage-sensitive determinant of senescence-associated phenotypes and DSB repair after IR. The same dosage sensitivity sets the two features that define the quality of the resulting senescent cell, namely its SASP amplitude and its apoptotic sensitivity. TIP60 levels do not affect the amount or timing of cell arrest after IR, but scale DSB repair kinetics. This is consistent with current knowledge of TIP60 at breaks. TIP60 is recruited to DSBs, where it acetylates histone H4 and H2AX ^62,72^, allowing flanking chromatin to be accessible, antagonizing 53BP1 retention, and promoting end resection and HR repair ^49,63^. If this activity depends on the level of TIP60 availability, TIP60-depleted cells are expected to repair slowly and retain persistent foci, while TIP60-overexpressing cells resolve them rapidly. Both 53BP1 and γH2AX foci behaved accordingly across the acute stress, transition and senescent phases, and the magnitude of the difference tracked the extent of the perturbation. The relationship between foci burden and proliferation reinforces this interpretation. The lines that completed the fewest population doublings carried the greatest number of persistent foci, the opposite of what a replication-dependent origin of those foci would predict. The differences in foci burden therefore reflect repair capacity, not residual cycling.

Despite these differences in repair, every line entered senescence. All lines reached a proliferation plateau with near-complete loss of DNA synthesis in the senescent phase, and SA-β-gal positivity was high in every TIP60 modulation. Depletion modestly raised the SA-β-gal-positive fraction, and overexpression left it unchanged. Depleted cells nonetheless secreted less IL-6 and expressed lower SASP transcript levels; both measured per cell. A larger senescent fraction would be expected to raise secretion, not lower it. The reduced SASP of TIP60-depleted cells therefore reflects the state of each senescent cell rather than the proportion of cells that became senescent.

Persistent DDR foci are often strongly associated with the SASP ^5,29^, although we have previously shown that chromatin stress can also associate with SASP activation in the absence of detectable DSBs ^28^. Much like chromatin stress, in the case of TIP60 manipulation, we observed a disconnection between DNA damage foci and secretion measured via IL-6 protein, SASP transcripts or the multiplex secretome. Persistent nuclear damage foci are therefore not sufficient to sustain a SASP when TIP60 is limiting. Micronuclei, by contrast, decreased with TIP60 depletion and increased with overexpression, tracking proliferation rather than foci burden. A dependence on cell division is expected, since micronuclei form only when damaged cells pass through mitosis ^64^. TIP60 also acts at both steps that convert cytosolic chromatin into secreted protein, since it acetylates cGAS to promote its activity^52^ and coactivates NF-κB-dependent transcription^68^. Our data therefore support a notion in which TIP60 controls the SASP in part through the cytosolic arm of the damage response rather than the nuclear one. It supplies both the substrate, in the form of micronuclei generated by residual division, and possibly the signalling competence to act on it. Depletion removes both, so an increased nuclear foci burden cannot compensate. The hypothesis remains to be tested. Scoring cGAS recruitment to micronuclei and NF-κB target occupancy across the series would test it directly, as would investigating whether cGAS or STING inhibition abolishes the differences between lines.

The apoptotic phenotype is attributed to the upstream difference in DNA damage repair. Slow repair and weak DDR signalling limit p53 output in the TIP60-depleted senescent cells. Reduced p53 output in TIP60-depleted lines lowered both its pro-apoptotic targets PUMA, NOXA, BAD and FAS and its arrest target p21, rather than selectively affecting the pro-apoptotic arm. Since TIP60-dependent acetylation of p53 favours pro-apoptotic over arrest targets^37,38^, such a non-selective reduction suggests that TIP60 loss limits p53 activity more broadly in senescent cells.

Spontaneous cell death during the transition and senescent phases decreased with TIP60 depletion and increased with TIP60 overexpression. ABT-263 showed reduced senolytic activity against TIP60-depleted cells and modest cytotoxicity towards TIP60-overexpressing cells relative to controls. The depleted lines were largely resistant to ABT-263, which may be explained by increased *MCL1* transcript levels. Whether TIP60 directly regulates MCL-1 remains unclear. Since ABT-263 selectively inhibits BCL-2, BCL-xL and BCL-w, MCL-1 upregulation confers resistance to BCL-2 family inhibitors^70,71^. TIP60-depleted senescent cells resist ABT-263 through both reduced pro-apoptotic p53 output and elevated MCL-1 expression. TIP60 level may therefore predict senolytic response and identify TIP60-low populations that require MCL-1 co-inhibition. These findings explore a broad problem, namely why cells receiving comparable damage adopt such different senescent fates and cell states. Our work identifies TIP60, a chromatin modifier, upstream of that divergence and indicates that a combination of repair capacity and apoptotic state, rather than lesion load alone, sets the future secretory and apoptotic state of the damaged cell. The uncoupling of nuclear foci burden from SASP amplitude further argues that persistent DDR signalling is not sufficient for a full SASP, when TIP60 is limiting. Micronucleus frequency tracked secretory output, pointing to a contribution from cytosolic DNA sensing that remains to be tested. Interestingly, from a translational point of view, if TIP60 level determines whether damaged cells are inflammatory and apoptosis-prone or quiet and drug-resistant, TIP60 status becomes a candidate biomarker for pro-senescence and senolytic treatment, with MCL-1 co-targeting a rational addition where TIP60 is low. Taken together, these results identify TIP60 dosage as a determinant of the senescence-apoptosis balance as well as the quality of senescent state for the point of view of apoptosis resistance and SASP. TIP60 therefore functions as a dosage-sensitive modifier of senescent cell identity, and its level may help predict which senescent populations respond to senolytic or senomorphic treatments.

## Acknowledgements

The Authors thank all the former and current members of the Rodier laboratory for valuable comments and discussions., We thank Isabelle Clement and the Institut du cancer de Montréal (ICM) Imaging and Live Imaging Platform. We thank all our funding agencies for their support: The team was supported by the Institut du cancer de Montréal, the professoral support program of the radiology, radio-oncology and nuclear medicine department of the faculty of medicine of the Université de Montréal, the Canadian Institute for Health Research (CIHR MOP114962 to F.R.), the Natural Sciences and Engineering Research Council of Canada (NSERC) (F.R.) and the Terry Fox Research Institute (TFRI) (F.R.). F.R. is a researcher at CRCHUM/ICM, which receives support from the FRQS. E.G. is supported by TFRI, OVCAN and ICM. N.M. is supported by TFRI and CHIR. S.D. is supported by TFRI and is a Fonds de Recherche du Québec Santé PhD fellow, ICM’s Prime Canderel Fellowship recipient, Department of Molecular Biology UdeM Excellence fellow and Cr-CHUM Excellence fellow.

## Authors’ contributions

S.D., N.M. and F.R. conceptualised the study. S.D. and N.M. designed the study, performed experiments and collected/analysed data. S.D., N.M., and F.R. wrote the manuscript. E.G., A.C.L., and G.B. provided technical support. F.R. supervised the work. F.R. provided financial support, expertise, conceptual advice, and revised the manuscript.

## Supplementary Figure Legends

**Figure S1. TIP60 dosage does not induce cell death or senescence in fibroblasts. Related to Figure 1**. (A, D) Representative live-cell images of (A) knockdown and (D) overexpression lines at days 1, 3, 5 and 7 after selection, acquired on an IncuCyte S3 system with a 10× objective. tGFP-NLS-labelled nuclei are shown in green and propidium iodide (PI)-positive dead cells in red, overlaid on phase contrast. (B, E) Quantification of cell death in the experiment shown in A and D for (B) knockdown and (E) overexpressing lines. PI was added to the medium at 0.5 µg/ml and cell death was calculated at each time point as the number of PI-positive nuclei relative to the total number of tGFP-NLS-positive nuclei. One representative of four independent experiments is shown: mean ± SD of 6 replicate wells. (C, F) Cumulative population doublings of (C) knockdown and (F) overexpressing lines in two further independent passaging experiments, each started at a different cumulative PD. (G, H) Percentage of SA-β-gal-positive cells in untreated knockdown (G) and overexpressing (H) lines at days 7, 14 and 21 after selection. Cells were stained for 16 h at pH 6.0, counterstained with DAPI, and the proportion of blue cells among DAPI-positive nuclei was determined in five randomly selected fields per well. Mean ± SD of three independent experiments. (I, J) Representative images of the SA-β-gal assay for knockdown (I) and overexpressing (J) lines at each time point, showing DAPI (blue) and SA-β-gal (brightfield). Images were acquired on an EVOS FL microscope with a 10× objective. Scale bar: 100 µm. B, C, E–H were analysed by two-way ANOVA followed by Dunnett’s multiple comparisons test against the respective control line (shGFP.3 or EGFP). *p < 0.05, **p < 0.05, ****p < 0.0005, ns: not significant.

**Figure S2. Representative images for the proliferation, DNA synthesis and DNA damage foci analyses after IR. Related to Figure 2**. (A) Knockdown and (B) overexpressing lines, representative real-time images on days 1, 3, 5, 7 and 9 post 10 Gy IR, showing GFP-positive nuclei (green) and PI-positive dead cells (red, 0.5 µg/ml). (C) Knockdown and (D) overexpressing lines, representative images of EdU incorporation (Click-iT Alexa Fluor 647) after 48 h labelling during the acute stress, transition and senescent phases (days 1–3, 5–7 and 7–9 post IR). Nuclei were counterstained with DAPI. (E) Knockdown and (F) overexpressing lines, representative immunofluorescence images of 53BP1 (red) and γH2AX (green) foci with DAPI counterstain (blue) and merged channels on days 3, 5 and 7 post IR. Insets show magnified views of the boxed regions. Scale bars: 100 µm and 20 µm (insets). Quantification is shown in Fig. 2C–J.

**Figure S3. TIP60 dosage modulates secretory output in non-irradiated cells, and representative SA-β-gal images after IR.** (A) Knockdown and (B) overexpressing lines, IL-6 secretion in non-irradiated cells measured by ELISA in CM collected at days 6–7, 13–14 and 20–21 after selection, after 16 h of incubation, expressed as fold change relative to controls at day 7. Statistical significance was assessed by two-way ANOVA with Dunnett’s multiple comparisons test against controls at each time point. (C) Knockdown and (D) overexpressing lines, expression of SASP genes measured by RT-qPCR in non-irradiated cells, shown as log2 fold change in (C) shTIP60.5 relative to shGFP.3 and in (D) TIP60HA relative to EGFP with −log10 p-value. Normalisation, statistics and thresholds as in Fig. 3C, 3D. (E) Knockdown and (F) overexpressing lines, secreted factors in CM from non-irradiated cells measured by multiplex immunoassay as in Fig. 3E, 3F, shown as log2 ratio relative to (E) shGFP.3 or (F) EGFP. C denotes non-irradiated control conditions. (G) Knockdown and (H) overexpressing lines, representative images of SA-β-gal staining (brightfield) with DAPI counterstain at day 10 post IR, 10× magnification. Scale bar, 100 µm. Quantification is shown in Fig. 3I, 3J. Data in A–F are one representative of three independent experiments, mean ± SD of 3 technical replicates. *p < 0.05, **p < 0.05, ****p < 0.0005, ns: not significant.

**Figure S4. TIP60 level does not affect cell death or ABT-263 sensitivity in non-irradiated cells**. (A) Knockdown and (B) overexpressing lines, cell death in non-irradiated cells treated with 1 µM ABT-263 at day 6 post seeding, measured every 4 h for 24 h. Representative images at 4, 8, 16 and 24 h post treatment show GFP-positive nuclei (green) and PI-positive dead cells (red). Data are one representative of three independent experiments, mean ± SD of 3 technical replicates. Statistical significance was assessed by two-way ANOVA with Dunnett’s multiple comparisons test against controls at each time point. ns not significant.

## Notes

### Competing Interest Statement

The authors have declared no competing interest.

