## Supplemental Figures and Tables for "TIP60/KAT5-mediated regulation of apoptotic sensitivity during DNA damage-induced senescence"

**Fig S1**

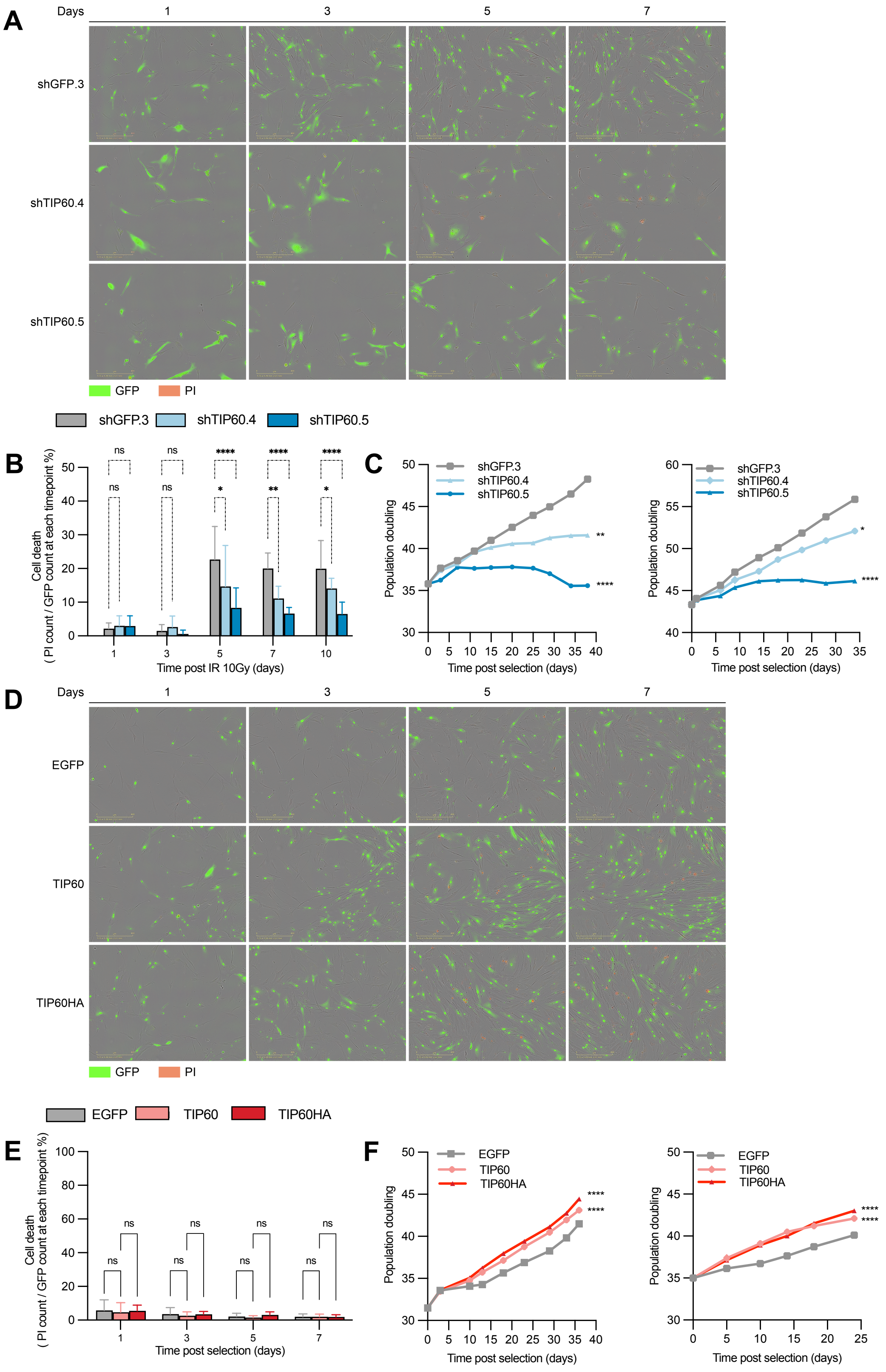
A
Days
1
3
5
7
shGFP.3
shTIP60.4
shTIP60.5
GFP
PI
shGFP.3
shTIP60.4
shTIP60.5
B
Cell death
( PI count / GFP count at each timepoint %)
Time post IR 10Gy (days)
ns
*
**
****
C
Population doubling
Time post selection (days)
D
Days
EGFP
TIP60
TIP60HA
GFP
PI
EGFP
TIP60
TIP60HA
E
Cell death
( PI count / GFP count at each timepoint %)
Time post selection (days)
F
Population doubling
Time post selection (days)

#### Fig S1

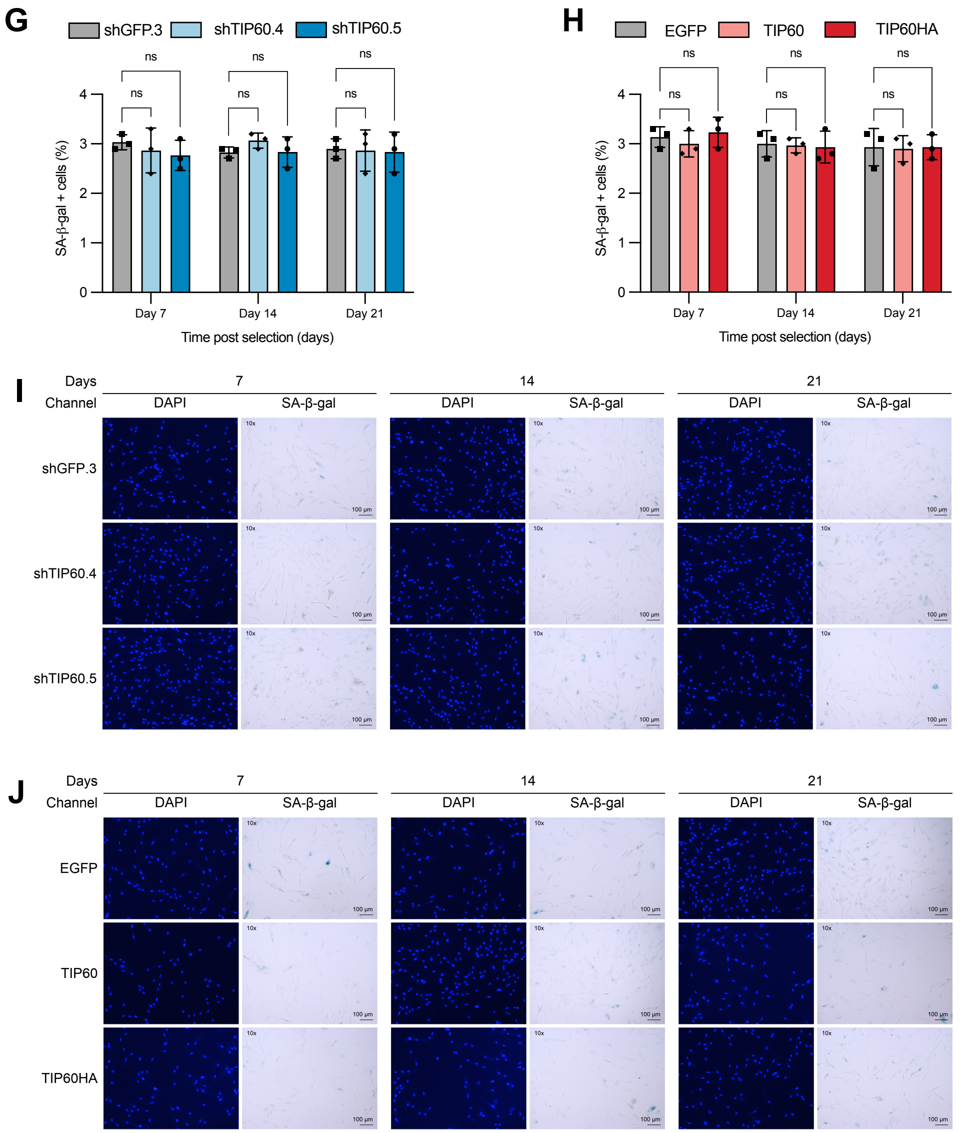

G
shGFP.3
shTIP60.4
shTIP60.5
ns
SA-β-gal + cells (%)
Day 7
Day 14
Day 21
Time post selection (days)
H
EGFP
TIP60
TIP60HA
ns
SA-β-gal + cells (%)
Day 7
Day 14
Day 21
Time post selection (days)
I
Days
7
14
21
Channel
DAPI
SA-β-gal
shGFP.3
shTIP60.4
shTIP60.5
J
Days
7
14
21
Channel
DAPI
SA-β-gal
EGFP
TIP60
TIP60HA

### Fig S2

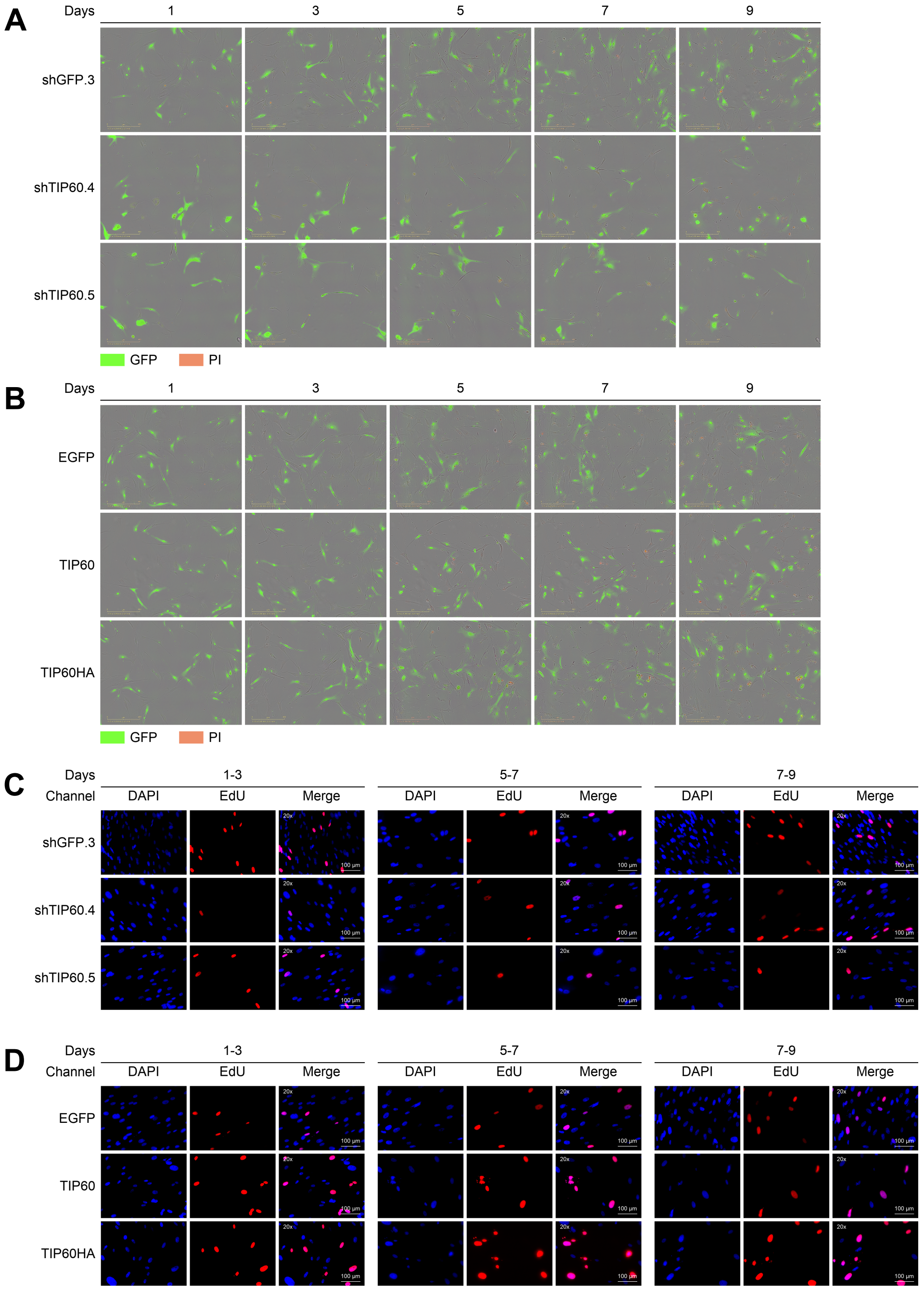
A
Days
1
3
5
7
9
shGFP.3
shTIP60.4
shTIP60.5
GFP
PI
B
Days
1
3
5
7
9
EGFP
TIP60
TIP60HA
GFP
PI
C
Days
1-3
5-7
7-9
Channel
DAPI
EdU
Merge
shGFP.3
shTIP60.4
shTIP60.5
20x
100 µm
D
Days
1-3
5-7
7-9
Channel
DAPI
EdU
Merge
EGFP
TIP60
TIP60HA
20x
100 µm

### Fig S2

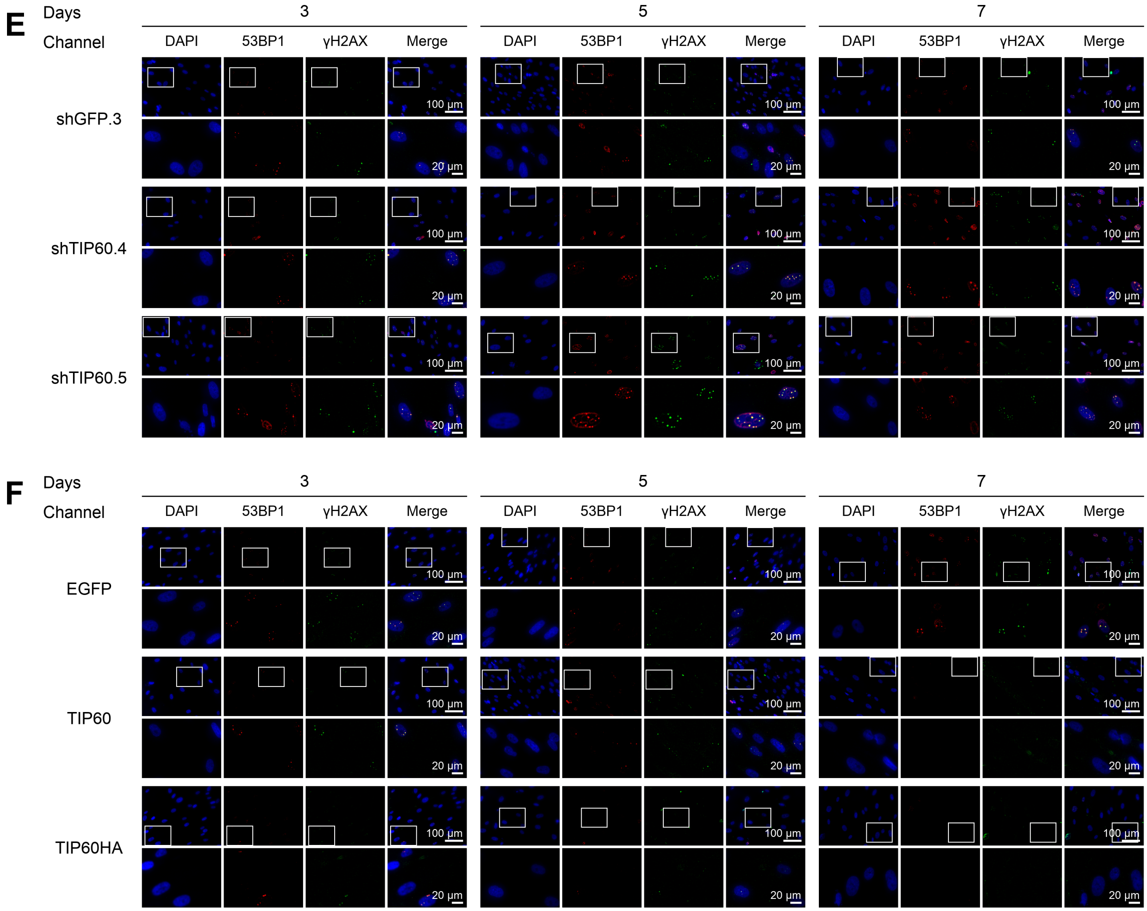
E
Days
3
5
7
Channel
DAPI
53BP1
γH2AX
Merge
shGFP.3
shTIP60.4
shTIP60.5
100 μm
20 μm
F
EGFP
TIP60
TIP60HA

#### Fig S3

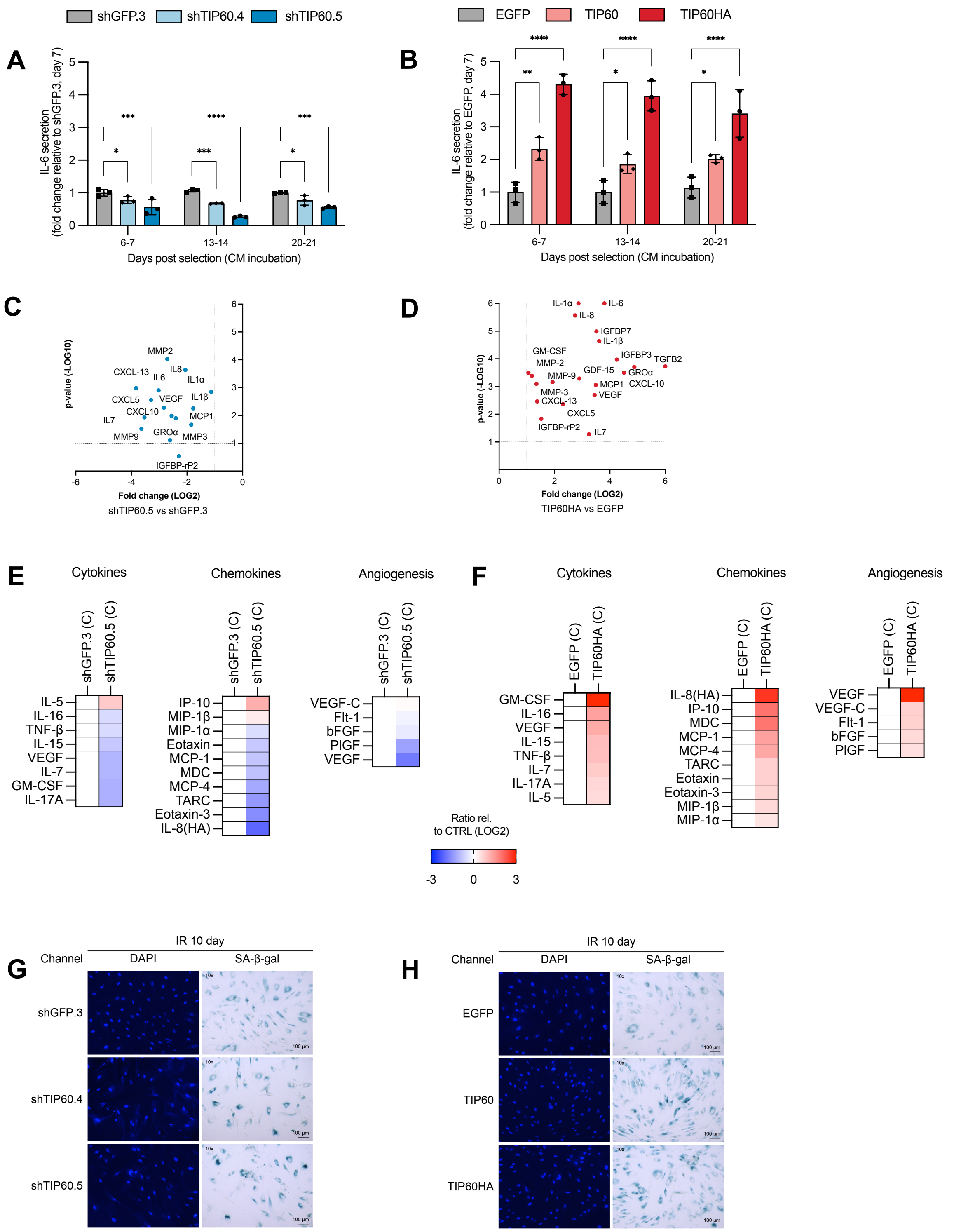

A
shGFP.3
shTIP60.4
shTIP60.5
IL-6 secretion (fold change relative to shGFP.3, day 7)
Days post selection (CM incubation)
6-7
13-14
20-21
B
EGFP
TIP60
TIP60HA
IL-6 secretion (fold change relative to EGFP, day 7)
Days post selection (CM incubation)
C
p-value (-LOG10)
Fold change (LOG2)
shTIP60.5 vs shGFP.3
MMP2
CXCL-13
IL8
IL6
IL1α
CXCL5
VEGF
IL1β
CXCL10
IL7
MCP1
MMP9
GROα
MMP3
IGFBP-rP2
D
p-value (-LOG10)
Fold change (LOG2)
TIP60HA vs EGFP
IL-1α
IL-6
IL-8
IGFBP7
IL-1β
GM-CSF
IGFBP3
TGFB2
MMP-2
GDF-15
GROα
MMP-9
MCP1
CXCL-10
MMP-3
VEGF
CXCL-13
CXCL5
IGFBP-rP2
IL7
E
Cytokines
Chemokines
Angiogenesis
shGFP.3 (C)
shTIP60.5 (C)
IL-5
IL-16
TNF-β
IL-15
VEGF
IL-7
GM-CSF
IL-17A
IP-10
MIP-1β
MIP-1α
Eotaxin
MCP-1
MDC
MCP-4
TARC
Eotaxin-3
IL-8(HA)
VEGF-C
Flt-1
bFGF
PIGF
VEGF
F
Cytokines
Chemokines
Angiogenesis
EGFP (C)
TIP60HA (C)
GM-CSF
IL-16
VEGF
IL-15
TNF-β
IL-7
IL-17A
IL-5
IL-8(HA)
IP-10
MDC
MCP-1
MCP-4
TARC
Eotaxin
Eotaxin-3
MIP-1β
MIP-1α
VEGF
VEGF-C
Flt-1
bFGF
PIGF
Ratio rel. to CTRL (LOG2)
-3
0
3
G
IR 10 day
Channel
DAPI
SA-β-gal
shGFP.3
shTIP60.4
shTIP60.5
H
IR 10 day
Channel
DAPI
SA-β-gal
EGFP
TIP60
TIP60HA

#### Fig S4

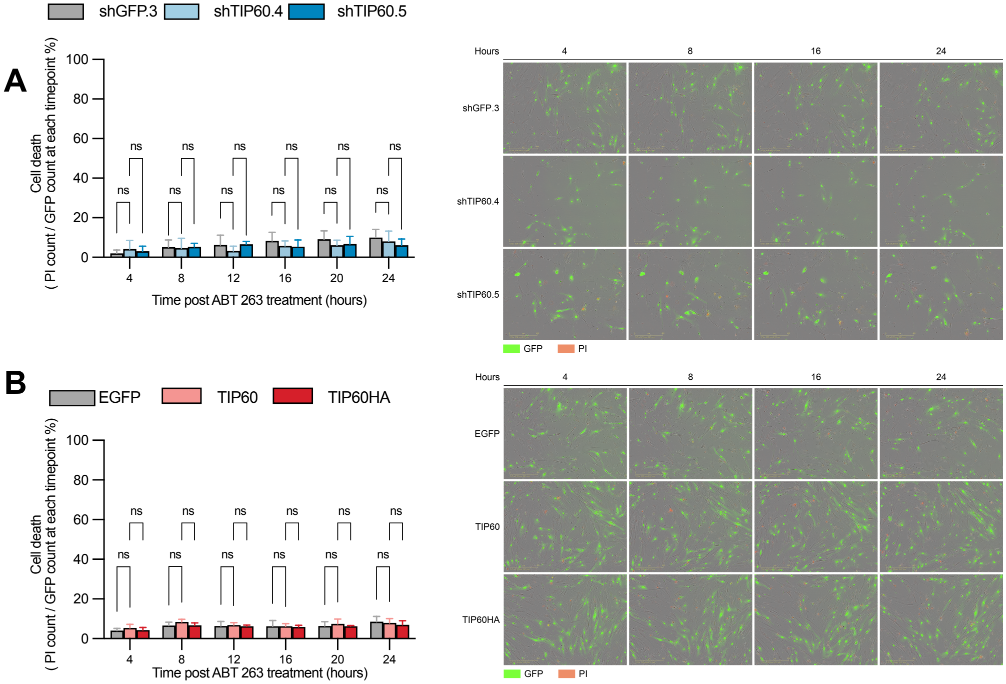

A
shGFP.3
shTIP60.4
shTIP60.5
Cell death
( PI count / GFP count at each timepoint %)
100
80
60
40
20
0
ns
4
8
12
16
20
24
Time post ABT 263 treatment (hours)
Hours
shGFP.3
shTIP60.4
shTIP60.5
GFP
PI
B
EGFP
TIP60
TIP60HA
Cell death
( PI count / GFP count at each timepoint %)
ns
Time post ABT 263 treatment (hours)
Hours
EGFP
TIP60
TIP60HA
GFP
PI

#### Table S1

| Construct | Target gene | Clone no. | Vector | Target sequence (5′→3′) | Selection |
|---|---|---|---|---|---|
| shTIP60.4 | KAT5 (TIP60) | TRCN0000020317 | pLKO.1-puro | CCTCCTATCCTATCGAAGCTA | Puromycin |
| shTIP60.5 | KAT5 (TIP60) | TRCN0000020318 | pLKO.1-puro | TCGAATTGTTTGGGCACTGAT | Puromycin |
| shGFP.3 | EGFP (control) | Addgene #30323 | pLKO.1-puro | GCAAGCTGACCCTGAAGTTCAT | Puromycin |

**Table S2**

| Gene | Forward 5′→3′ | Reverse 5′→3′ |
|---|---|---|
| TBP | GAGCCAAGAGTGAAGAACAG | ACCTTATAGGAAACTTCACATCAC |
| KAT5 (TIP60) | CACAGATCACCATCAATGAGATTA | CTCTGACAGTGTGAGGATGTA |
| BAX | TTTCTGACGGCAACTTCAA | TGGTGCACAGGGCCTTGA |
| BAD | CCAACCTCTGGGCAGCACAGC | TTTGCCGCATCTGCGTTGCTGT |
| BCL2 | AACATCGCCCTGTGGATGAC | GGCCGTACAGTTCCACAAAG |
| MCL1 | GGACATCAAAAACGAAGACG | GCAGCTTTCTTGGTTTATGG |
| PUMA | ACGACCTCAACGCACAGTACGA | CCTAATTGGGCTCCATCTCGGG |
| NOXA | CTGGAAGTCGAGTGTGCTACTC | TGAAGGAGTCCCCTCATGCAAG |
| p53 (TP53) | GGAAGACTCCAGTGGTAATCTA | TTGGGCAGTGCTCGCTTA |
| p21 (CDKN1A) | GGGACAGCAGAGGAAGAC | TGGAGTGGTAGAAATCTGTCA |
| FAS | GGACCCAGAATACCAAGTGCAG | GTTGCTGGTGAGTGTGCATTCC |
| NOTCH1 | GGTGAACTGCTCTGAGGAGATC | GGATTGCAGTCGTCCACGTTGA |
| GDF15 | GCCAGAAGTGCGGCTGGGA | GGACAGCCGGAACAGAGCC |
| IL6 | TGTGTGAAAGCAGCAAAGA | GGCAAGTCTCCTCATTGAA |
| IL1A | TGTATGTGACTGCCCAAGATGAAG | AGAGGAGGTTGGTCTCACTACC |
| IL1B | CCACAGACCTTCCAGGAGAATG | GTGCAGTTCAGTGATCGTACAGG |
| IL7 | ATGAGAGTGTTCTAATGGTC | GGAGCATTCAGTTTCCATTG |
| IL8 (CXCL8) | GCCAACACAGAAATTATTGTAAAG | TTATGAATTCTCAGCCCTCTTC |
| CXCL5 | CAGACCACGCAAGGAGTTCATC | TTCCTTCCCGTTCTTCAGGGAG |
| CXCL10 | GGTGAGAAGAGATGTCTGAATCC | GTCCATCCTTGGAAGCACTGCA |
| CXCL13 | TATCCCTAGACGCTTCATTGATCG | CCATTCAGCTTGAGGGTCCACA |
| CXCL1 (GROα) | CCAAGAACATCCAAAGTGTGAAC | AGTTGGATTTGTCACTGTTCA |
| CCL2 (MCP1) | GCTGTGATCTTCAAGACCAT | GTTTGCTTGTCCAGGTGG |
| MMP2 | AGCGAGTGGATGCCGCCTTTAA | CATTCCAGGCATCTGCGATGAG |
| MMP3 | CACTCACAGACCTGACTCGGTT | AAGCAGGATCACAGTTGGCTGG |
| MMP9 | GCAGCTGGCAGAGGAATACC | TTTGGAATCTGCCCAGGTCT |
| VEGFA | TTGCCTTGCTGCTCTACCTCCA | GATGGCAGTAGCTGCGCTGATA |
| CSF2 (GM-CSF) | GGAGCATGTGAATGCCATCCAG | CTGGAGGTCAAACATTTCTGAGAT |
| IGFBP3 | AACGCTAGTGCCGTCAGC | CGGTCTTCCTCCGACTCAC |
| IGFBP7 | GGCCCAGAAAAGCATGAAG | TCATATTCTCCAGCATCTTCCTTA |
| IGFBP-rP2 (CTGF) | CCTGCAGGCTAGAGAAGCAGA | TGCACTTTTTGCCCTTCTTAATGT |
| TGFB2 | CCAAAGGGTACAATGCCAAC | CAGATGCTTCTGGATTTATGGTATT |
